# Distinct yet complementary roles of temporal and spectral predictions in auditory detection

**DOI:** 10.64898/2026.03.04.709562

**Authors:** Johannes Wetekam, Chloé Dumeige, Manon Beurtey, Sophie K. Herbst

## Abstract

Predictive processing enables the brain to anticipate both *when* and *what* sensory events will occur. Although temporal and spectral predictions are both known to facilitate perception, their functional contributions and interaction remain poorly understood, particularly in uncertain listening conditions where predictions must be inferred from implicit statistical regularities rather than explicit cues or rhythms. Here, we orthogonalized temporal and spectral predictability in a non-rhythmic auditory detection task and used a signal-detection framework to dissociate perceptual and decisional processes in humans of either sex. Temporal predictions primarily increased response readiness: predictable target timing led to faster responses and more hits but also more false alarms, reflecting a liberal shift in response bias. Spectral predictions, in contrast, selectively enhanced perceptual sensitivity without a comparable speeding of responses. Crucially, despite their distinct behavioral effects, temporal and spectral predictions interacted synergistically, such that perceptual sensitivity was greatest when both sound timing and frequency were predictable, consistent with a spectrotemporal filter mechanism. Furthermore, temporal and spectral variability were encoded differently: performance adapted strongly to the distribution of foreperiod durations but remained largely stable across the spectral distribution, suggesting that the brain exploits temporal and spectral statistics in distinct ways when forming predictions under uncertainty. Together, these findings demonstrate that temporal and spectral predictions make complementary contributions to auditory decision-making, interact synergistically to improve perceptual sensitivity, and rely on distinct encoding strategies for temporal and spectral variability.

**Significance statement:** In everyday listening, the brain must predict both *when* relevant sounds will occur and *what* they will sound like. Although temporal (“*when*”) and spectral (“*what*”) predictions are both known to support perception, their distinct functions and possible synergies are not well understood. Here, using a non-rhythmic auditory detection task and a signal-detection framework, we show that temporal predictions mainly increase response readiness, whereas spectral predictions selectively enhance perceptual sensitivity. When both are available, they interact synergistically to maximize sensitivity. These findings reveal that temporal and spectral predictions make complementary contributions to auditory decision-making and show how the brain combines multiple predictive dimensions to improve perception under uncertainty.

## 1. Introduction

Detecting behaviorally relevant sounds under acoustic uncertainty is a central challenge for the auditory system. In natural environments, sounds are embedded in noise, occur at irregular times, and vary in their spectral content. One way the auditory system can reduce this uncertainty is by exploiting regularities in the sensory input to generate predictions about forthcoming events. In audition, stimulus timing (“when”) and stimulus content (“what”) are two particularly important sources of predictive information. A substantial body of work has shown that temporal and spectral predictions can each facilitate auditory processing (for reviews see Arnal and Giraud, 2012; Heilbron and Chait, 2018; Nobre and van Ede, 2018).

Spectral (or content) predictions have been shown to improve behavioral performance (Lange, 2008; Morillon et al., 2016; Wollman and Morillon, 2018) and to modulate auditory responses to predicted stimuli (Schwartze et al., 2013; Carbajal and Malmierca, 2018; Demarchi et al., 2019). Temporal predictions have been associated with motor preparation (Morillon et al., 2015; Auksztulewicz et al., 2018; Rimmele et al., 2018), a speeding of responses in sensory decision-making tasks (Lange, 2008; Hsu et al., 2013; Morillon et al., 2016), as well as improved spectral processing, such as pitch discrimination (Herbst and Obleser, 2017, 2019; Chang et al., 2019; Bonnet et al., 2024). Thus, both “when” and “what” predictions can shape auditory behavior, but their distinct functional contributions remain difficult to separate. On the one hand, temporal predictability has been linked to stimulus-unspecific gain modulations and increased response readiness, suggesting a broad preparatory mechanism that facilitates processing at expected moments in time (Auksztulewicz et al., 2018, 2019). On the other hand, several studies indicate that temporal prediction benefits can depend on the availability of spectral predictions (Hsu et al., 2013; Morillon et al., 2016; Wollman and Morillon, 2018), consistent with a spectrotemporal filtering mechanism in which temporal predictions selectively enhance processing of behaviorally relevant sensory channels (Lakatos et al., 2013). These findings show that temporal and spectral predictions are closely linked but leave unresolved how each dimension specifically contributes to auditory perception and how they interact when both are present.

Here, we aim to address this issue by orthogonalizing temporal and spectral predictability in a non-rhythmic auditory detection task (Fig. 1a). In both dimensions, predictions had to be formed from implicit probability distributions of target timing (foreperiods) and frequency, respectively (Fig. 1b). We then used a signal detection theory (SDT)-based decomposition of participants’ performance to separate prediction effects on hit rates (HR), false alarms (FA), perceptual sensitivity, and response bias. Perceptual sensitivity describes how well participants can distinguish signal from noise, whereas response bias describes their tendency to report that a signal was present, independent of sensitivity (Green and Swets, 1966; Summerfield and de Lange, 2014). This approach allowed us to test whether temporal and spectral predictions influence the same or different components of detection behavior and whether their benefits interact such that perceptual sensitivity is greatest when both “when” and “what” are predictable, as would be expected from a spectrotemporal filter (Fig. 1c). Because forming predictions depends on exploiting temporal and spectral regularities, we finally examined how performance adapted to the statistical structure of each dimension by modelling behavior as a function of foreperiod duration and target frequency.

**Figure 1-.**
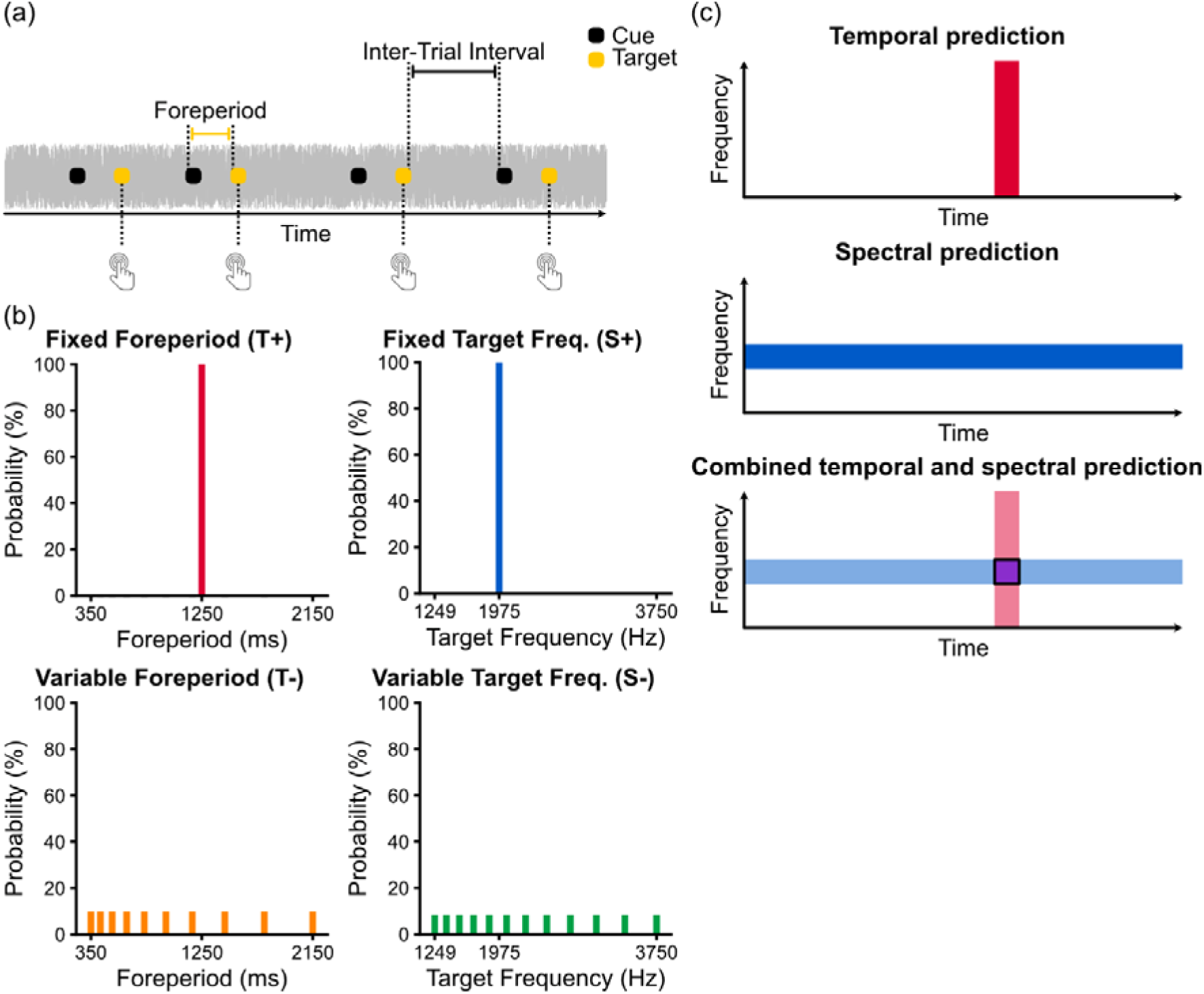
Foreperiod paradigm, predictability conditions, and spectrotemporal filter. (a) Schematic illustration of the foreperiod paradigm used in this study. Cue-target tone pairs were presented against continuous background noise. The time interval between the cue and target defines the foreperiod. Inter-trial intervals were jittered to prevent the emergence of isochronous stimulation patterns. In 10 % of trials, the target was omitted (catch trials). (b) Probability distributions of foreperiods and target frequencies. Temporal predictability was manipulated by keeping the foreperiod either fixed (top-left, red; T+) or variable (bottom-left, orange; T−) within a block. Spectral predictability was manipulated by keeping the target frequency either fixed (top-right, blue; S+) or variable (bottom-right, green; S−). Temporal and spectral predictability manipulations were crossed, leading to four different conditions: T+S+, T+S-, T-S+, T-S-. (c) Conceptual illustration of how temporal predictions (red) may constrain processing in time and spectral predictions (blue) may constrain processing in frequency, with their intersection (magenta) defining a spectrotemporal filter.

## 2. Material and Methods

### Participants

For this study, a total of 41 participants aged between 23 and 53 years (27 females, 14 males; mean age = 35.5 years, standard deviation (SD) = 7.8 years) were recruited via online announcements distributed through the PRISME platform mailing list (Institut du Cerveau, Paris). Exclusion criteria comprised self-reported auditory impairments and color blindness. All participants provided written informed consent prior to participation and received financial compensation (20 € in cash per session). The study was approved by the local ethics committee of Université Paris-Saclay (Reference: CER-Paris-Saclay-2022-004/A3) and was conducted in accordance with the World Medical Association (WMA) Declaration of Helsinki.

### Stimuli and data acquisition

All experiments were conducted at the Paris Brain Institute PRISME Core Facility (RRID:SCR_026394), Paris, France. A classroom equipped with 12 individual computers enabled parallel data acquisition within a single session. Stimulus presentation, visual display, and behavioral data collection were implemented using the Python library Expyriment (Krause and Lindemann, 2014). Auditory stimuli were delivered through Logitech G35 headphones at a sampling rate of 44100 Hz.

The experiment was based on a foreperiod paradigm combined with an auditory detection task (Fig. 1a). Sound sequences were presented against continuous background noise. The background noise consisted of low-pass filtered white noise (cut-off frequency: 5 kHz) and was continuously played throughout each block. Each trial consisted of a pair of tones, with the first tone serving as a cue and the second tone as a target sound. Cue tones had a duration of 250 ms and a fixed frequency of 1975 Hz. Target tones had a duration of 50 ms, and their frequency depended on the experimental condition (see below). All tones were pure sine waves and included 10 ms cosine-shaped rise and fall ramps.

Prior to the foreperiod task, the intensity of both the background noise and the tones was individually determined for each participant using staircase-based procedures. First, the background noise level was adjusted relative to each participant’s individual sensation level using an ascending and descending method of limits. The final noise level was fixed at 40 dB above the individual sensation level. Next, hearing thresholds for pure tones were measured within this background noise using an adaptive staircase procedure. A 2-down-1-up rule was applied, such that tone intensity was decreased after two consecutive detections and increased after a single miss. This procedure converges on a detection level of approximately 70 % correct responses. Thresholds were estimated separately for three tone frequencies (1250, 1975, and 3750 Hz) and linearly interpolated for the remaining frequencies used in the experiment. During the foreperiod task, cue tones were presented at a level 20 dB above the individual hearing threshold, whereas target tones were presented at hearing threshold level within the background noise. This resulted in clearly audible cues followed by difficult-to-detect target tones.

In 10 % of the trials, the cue was not followed by a target sound (catch trials). This proportion of catch trials was chosen to allow the estimation of false alarm rates, while keeping a high number of signal trials so that participants had sufficient exposure to the foreperiod and target-frequency distributions to form strong predictions. Participants were instructed to press the spacebar as quickly as possible whenever they detected a target tone, while avoiding responses on trials in which no target tone was presented. Responses made before target onset were classified as premature responses. Following target onset, participants had a response window of 1 s. After each tone pair, visual feedback indicating the response outcome (too early, hit, miss, correct rejection, or false alarm) was presented on the screen. Feedback was displayed immediately following the participant’s response or, in the absence of a response, after the 1 s response window had elapsed. A short training sequence of 40 trials preceded the main experimental session.

Temporal and spectral predictability were manipulated using a 2 × 2 block wise design. Temporal predictability was varied by keeping the foreperiod duration either fixed at 1250 ms within a block (high temporal predictability, T+) or randomly drawing it from a log-spaced uniform distribution ranging from 350 to 2150 ms (350, 428, 523, 641, 784, 959, 1173, 1436, 1757, and 2150 ms; low temporal predictability, T-). Spectral predictability was manipulated via the target frequency, which was either fixed at 1975 Hz within a block (high spectral predictability, S+) or randomly drawn from a log-spaced uniform distribution ranging from 1249 to 3750 Hz (1249, 1381, 1526, 1686, 1863, 2059, 2275, 2514, 2779, 3071, 3393, and 3750 Hz; low spectral predictability, S-; Fig. 1b). The fixed foreperiod and target-frequency values were not positioned equivalently within their respective variable ranges. The fixed foreperiod was located near the arithmetic mean of the foreperiod range, whereas the fixed target frequency was located close to the logarithmic mean of the frequency range. Crossing spectral predictability (S+ / S-) with temporal predictability (T+ / T−) yielded four experimental conditions (T+S+, T+S-, T-S+, and T-S-), one of which was presented per block. Each condition was presented in three blocks of 100 trials, yielding a total of 12 blocks (1200 trials) per participant. Block order was randomized across participants. Trials were separated by a jittered inter-trial interval drawn from a truncated normal distribution (mean = 1375 ms, SD = 62.5 ms, range = 1250–1500 ms), resulting in a non-isochronous stimulation pattern. Crucially, participants were not explicitly informed about the predictability manipulations. Their only instruction was to report the occurrence of target sounds as quickly and accurately as possible. The experimental task lasted approximately 1 h. Including instructions, training, and breaks, the full session lasted approximately 1.5 h.

### Data processing and statistical evaluation

Data analysis was conducted using custom scripts written in Python and R.

Behavioral performance was quantified using five dependent variables derived from participants’ key presses: reaction time (RT), hit rate (HR), false alarm rate (FA), sensitivity, and response bias. Reaction time was defined as the latency between target onset and the key press and was evaluated on detected signal trials (hits) only. Hit rate was defined as the probability of responding on signal trials and false alarm rate as the probability of responding on catch trials. Sensitivity and response bias were derived from hit rate and false alarm rate.

Statistical analyses were conducted using mixed-effects models to account for the repeated-measures structure of the data. Because reaction times are a continuous outcome, we analyzed them using a linear mixed-effects model (LMM). Response occurrence was analyzed using a binomial generalized linear mixed-effects model (GLMM), as each trial resulted in a binary outcome indicating whether or not a response was made. Hit rate, false alarm rate, sensitivity, and response bias were derived from this model. All models were implemented using the *lme4* package in R (Bates et al., 2015).

Sensitivity was defined as the exponentiated difference between the log odds of responding on signal and catch trials:

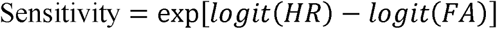

A value of 1 indicates that responses were equally likely on signal and catch trials. Values above 1 indicate that responses were more likely on signal than on catch trials, with larger values reflecting greater sensitivity. Response bias was defined as the exponentiated average of the log odds of responding on signal and catch trials:

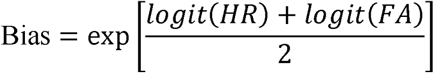

A value of 1 indicates a neutral response bias. Values above 1 indicate a liberal response bias, whereas values below 1 indicate a conservative response bias. Unlike conventional *d’* and *criterion* measures, sensitivity and response bias could be derived directly from the fitted mixed-effects model. This allowed main effects, interactions, and simple effects to be estimated and statistically tested within a single modelling framework, while accounting for the different numbers of signal and catch trials.

For participant-level descriptive values and Bayesian analyses, hit and false alarm rates were log-linearly corrected according to Hautus (1995):

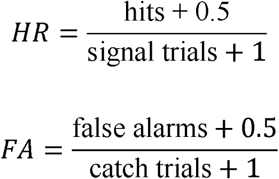

This correction ensured that participant-level probabilities remained strictly between 0 and 1.

Exclusion criteria were applied at both the participant and trial level. At the participant level, participants were excluded if they did not complete the full experiment (fewer than 12 blocks; *n* = 3). From the remaining sample, participants with overall hit rates below 40 % or above 95 % were excluded because their performance strongly deviated from the detection level of approximately 70 % targeted by the staircase procedure and therefore suggested unsuccessful calibration or insufficient task engagement. Three participants were excluded for hit rates below 40 %, and one participant was excluded for hit rates above 95 %. These exclusions resulted in a final sample of 34 participants included in the statistical analysis. At the trial level, premature responses (key presses occurring before target onset) were identified and labelled as “too early” and excluded from all analyses. In addition, hit trials with reaction times shorter than 100 ms were removed, as such responses are typically considered biologically implausible. Because temporal predictability could in principle lead to very rapid responses, all analyses were repeated without applying this reaction-time exclusion criterion to ensure that it did not affect the reported results. This robustness check did not change the pattern or statistical significance of the findings.

To assess the main effects of temporal (T) and spectral predictability (S), as well as their interaction and associated simple effects, RTs were analyzed using an LMM fitted to correct responses (hits) on signal trials only. The model included fixed effects of T, S, and their interaction (T×S), along with random intercepts and random slopes for T and S at the participant level. Response-based measures (hit rate, false alarm rate, sensitivity, and response bias) were analyzed using a GLMM. Hit rate and false alarm rate were modelled jointly within a single framework by including signal presence (signal vs. catch trials), temporal predictability (T), spectral predictability (S), and their interactions as fixed effects. This approach enabled the simultaneous estimation of hit and false alarm responses and the derivation of sensitivity and response bias from the same model. The GLMM included random intercepts and random slopes for signal presence, temporal predictability, and spectral predictability at the participant level. Planned contrasts were used to estimate main effects, interaction effects, and simple effects, with p-values adjusted for multiple comparisons using false discovery rate (FDR) correction (Benjamini and Hochberg, 1995). Corrected p-values lower than 0.05 were considered significant.

For non-significant effects relevant for data interpretation, additional Bayesian one-sample t-tests on participant-level were conducted. The tests were conducted using the *BayesFactor* package in R (Morey et al., 2026). Bayes factors are reported as BF_01_, with values above 1 indicating that the data were more likely under the null hypothesis (no effect) than under the alternative hypothesis, and values above 3 interpreted as moderate evidence in favor of the null hypothesis.

A second set of analyses examined how performance varied across the distributions of foreperiod durations and target frequencies. In this analysis, performance was quantified using reaction time and hit rate only. Sensitivity, response bias, and false alarm rate were not analyzed in this context, as catch trials do not provide information about foreperiod duration or target frequency. Reaction time was analyzed using an LMM, and hit rate was analyzed using a GLMM, with foreperiod duration or target frequency entered as continuous predictors modelled using second-order, mutually orthogonal polynomials. Temporal and spectral predictability were included as fixed effects in the models, and random intercepts were included at the participant level. Random slopes were not included, as models with random slopes did not converge and provided no improvement in model fit. For the model estimates, the predictability factor not directly related to the distribution of interest (i.e. spectral predictability for foreperiod analyses and temporal predictability for target-frequency analyses) was collapsed across its two levels. To quantify the effect size of the curvature of performance across the variable foreperiod and target frequency distributions, a curvature effect size was computed as the difference between the model-predicted minimum and maximum performance values across each distribution. The direction of this effect was defined such that improved performance around the distribution midpoint relative to the edges resulted in a positive effect (or an odds ratio (OR) greater than 1), whereas worse performance at the midpoint relative to the edges resulted in a negative effect (or an OR smaller than 1).

To further characterize how performance depends on the foreperiod distribution, we compared candidate temporal-prediction models based on the probability density function (PDF) and hazard-rate of the foreperiod distribution. Following Grabenhorst et al. (2019, 2026), we computed blurred PDFs by replacing each possible foreperiod with a Gaussian kernel. We tested two forms of blurring: temporal and constant blurring. For temporal blurring, the Gaussian width of each kernel was set to 21 % of the respective foreperiod, resulting in wider Gaussian widths at longer foreperiods. In contrast, for constant blurring, the same Gaussian width was used for all foreperiods. This constant width was set to the average of the widths at the shortest and longest foreperiods, resulting in 0.2625 s. Both the blurring procedure and the 21% scaling factor followed Grabenhorst et al. (2019, 2026). Because all foreperiods in our variable condition had the same probability, our constantly-blurred model corresponds to the probabilistic-blurring model in Grabenhorst et al. (2019, 2026). Hazard-rate predictors were derived either directly from the discrete foreperiod distribution or from the corresponding blurred PDFs. Because 10 % of trials were catch trials, hazard-rates were adjusted for the probability that no target would occur. Following the main model comparison in Grabenhorst et al. (2019, 2026), PDF predictors were entered as reciprocal PDF values, and hazard-rate predictors were mirrored. The mirrored hazard-rate captures the classical assumption of a linear inverse relationship between hazard-rate and reaction time, whereas the reciprocal PDF captures a non-linear inverse relationship between event probability and reaction time. Thus, larger predictor values indicated weaker temporal expectation. Each predictor was entered as a single fixed effect in a mixed-effects model with a random intercept for participant. Reaction times were analyzed with an LMM on detected target trials, and hit rate was analyzed with a binomial GLMM on signal trials. Models were compared using AIC.

### Pilot experiment with narrower frequency range and no catch trials

Prior to the main experiment, we conducted an independent pilot experiment with another 38 participants aged between 18 and 45 years (25 females, 13 males; mean age = 28.1 years, SD = 7.2 years) to assess behavioral effects of temporal and spectral predictability. The pilot experiment was identical to the main experiment in terms of recruitment procedures, exclusion criteria, task structure, stimulus timing, and analysis approach. The key differences were that the pilot used a substantially narrower target-frequency range (1724-2224 Hz) in the variable spectral condition and did not include catch trials. The absence of catch trials prevented the calculation of sensitivity and response bias and resulted in generally higher hit rates, likely leading to ceiling effects. For this reason, the pilot experiment was not suitable for addressing the main questions of the present study and motivated the design of the main experiment. Here, we use the pilot data only to assess whether performance across variable foreperiod and target-frequency distributions shows a similar qualitative pattern when the spectral range is reduced. Pilot data were analyzed using the same mixed-effects modelling approach as in the main experiment, focusing on how hit rate and reaction time varied as a function of foreperiod duration and target frequency under variable predictability. Results of the pilot experiment are reported in Supplementary Figure 1 and are referenced in the Discussion solely to evaluate the robustness of the distribution-level effects.

## 3. Results

### Temporal and spectral predictions play complementary roles in auditory detection

The current study examined the effects of temporal and spectral predictions on different aspects of auditory performance: hit rate, false alarm rate, reaction time, sensitivity, and response bias.

**Hit rate**: Analysis of hit rate revealed a significant main effect of temporal predictability, as well as a significant interaction between temporal and spectral predictability (T: p < 0.001; S: p = 0.055; T×S: p < 0.001; Fig. 2a-b). This interaction reflected a larger increase in hit rate due to temporal predictions when spectral predictions were present than when they were absent (T at S- and S+: p < 0.001). Spectral predictions did not significantly affect hit rate in the absence of temporal predictions but increased hit rate when temporal predictions were present (S at T-: p = 0.811, BF_01_ = 5.44; S at T+: p = 0.0018; Fig. 2c).

**Figure 2-.**
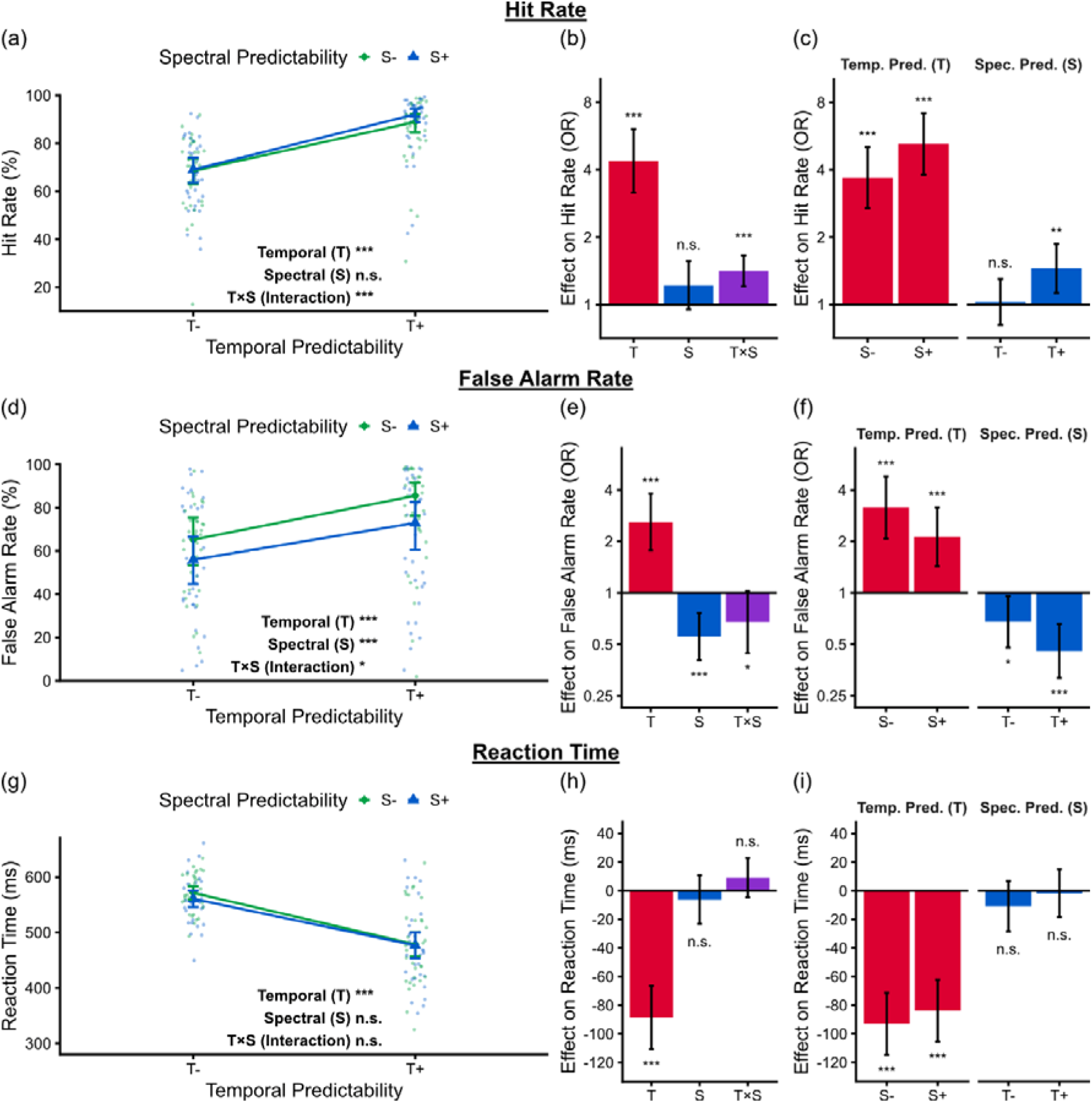
Effects of temporal and spectral predictability on behavioral performance. (a) Hit rate (%) as a function of temporal predictability (x-axis) and spectral predictability (color-coded). Symbols show model-estimated marginal means for each condition; whiskers indicate 95 % confidence intervals. Small dots represent individual participant means. (b) Main effects of temporal (T) and spectral (S) predictability, as well as their interaction (T×S), on hit rate, expressed as odds ratios (OR) derived from the GLMM. (c) Simple effects of temporal and spectral predictability on hit rate (OR), illustrating how the effect of one predictability factor depends on the level of the other. (d–f) Same as panels (a–c), but for false alarm rate. (g) Reaction time (ms) as a function of temporal and spectral predictability. (h) Main effects of temporal and spectral predictability, as well as their interaction, on reaction time, derived from the LMM. (i) Simple effects of temporal and spectral predictability on reaction time.

**False alarm rate**: False alarm rate was significantly modulated by temporal predictability, spectral predictability, and their interaction (T: p < 0.001; S: p < 0.001; T×S: p = 0.024; Fig. 2d-e). Importantly, the two prediction types had opposite effects: temporal predictions increased false alarm rate, whereas spectral predictions reduced it. In addition, a significant interaction indicated a reduction in false alarm rate when both prediction types were present together compared to when they occurred separately (T at S- and S+: p < 0.001; S at T-: p = 0.012; S at T+: p < 0.001; Fig. 2e-f).

**Reaction time**: In contrast to the response-based measures described above, reaction time was significantly reduced by temporal predictability, whereas spectral predictability and the temporal-spectral interaction had no significant effect (T: p < 0.001; S: p = 0.3786, BF_01_ = 3.10; T×S: p = 0.1637, BF_01_ = 3.55; Fig. 2g-i).

**Sensitivity**: Sensitivity, which integrates changes in both hit rate and false alarm rate, was significantly affected by temporal predictability, spectral predictability, and their interaction (T, S, and T×S: p < 0.001; Fig. 3a-b). Simple effects analyses revealed that temporal predictions improved sensitivity when spectral predictions were present, whereas the effect was not significant when spectral predictions were absent (T at S+: p < 0.001; T at S-: p = 0.2636, BF_01_ = 2.48). By contrast, spectral predictions also improved sensitivity when present in isolation, although this effect was weaker than when temporal predictions were available as well (S at T- and T+: p < 0.001; Fig. 3c).

**Figure 3-.**
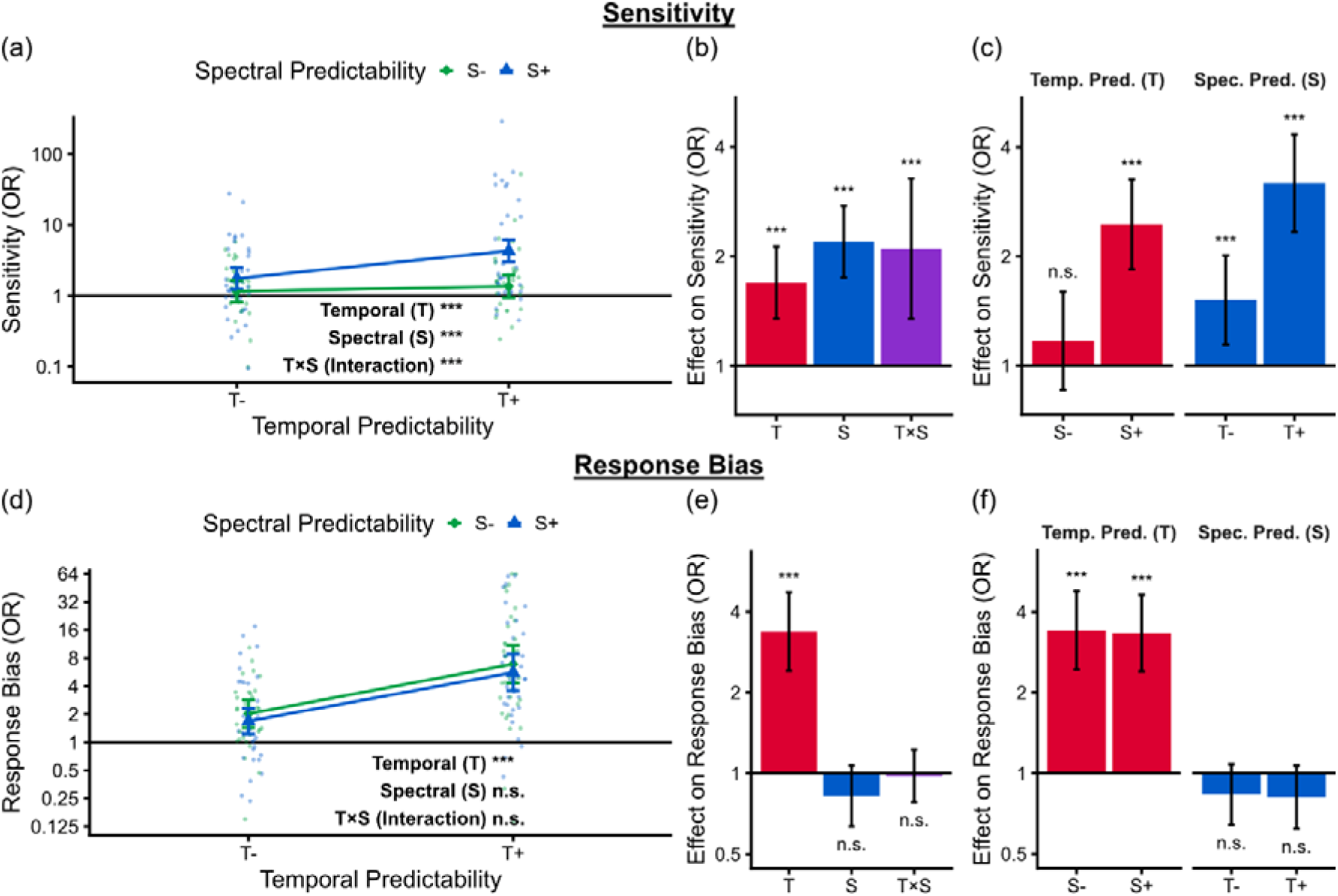
Effects of temporal and spectral predictability on sensitivity and response bias. (a) Sensitivity, expressed as odds ratios (OR) derived from the GLMM, as a function of temporal predictability (x-axis) and spectral predictability (color-coded). Symbols show model-estimated marginal means for each condition; whiskers indicate 95 % confidence intervals. Small dots represent individual participant means. (b) Main effects of temporal (T) and spectral (S) predictability, as well as their interaction (T×S), on sensitivity (OR). (c) Simple effects of temporal and spectral predictability on sensitivity (OR), illustrating how the effect of one predictability factor depends on the level of the other. (d–f) Same as panels (a–c), but for response bias.

**Response bias:** Temporal predictability significantly shifted response bias in a more liberal direction, whereas spectral predictability and the spectrotemporal interaction had no significant effect (T: p < 0.001; S: p = 0.1114, BF_01_ = 3.69; T×S: p = 0.8020, BF_01_ = 5.41; Fig. 3d–f).

### Distinct encoding strategies for temporal and spectral distributions

The dissociable effects of temporal and spectral predictions motivated a closer examination of how observers encode and exploit variability along these two dimensions. Performance under high variability can reveal whether, and in what way, observers encode and adapt to the statistical properties of each distribution. Therefore, we next examined how performance varied across foreperiods and target frequencies in the variable conditions. Specifically, we analyzed hit rate and reaction time as a function of foreperiod duration or target frequency, respectively, using a GLMM for hit rate and an LMM for reaction time. In these models, foreperiod duration or target frequency was entered as a continuous predictor, allowing the assessment of non-linear changes in performance across each distribution. Hit rate exhibited a pronounced inverse U-shaped relationship with foreperiod duration in the variable temporal condition (Fig. 4a). Notably, the modulation was slightly asymmetric, with peak performance shifted towards longer intermediate foreperiods rather than being centered exactly on the distribution midpoint. The U-shape pattern was supported by a significant quadratic term in the GLMM and reflected higher performance around the midpoint of the foreperiod distribution compared to its edges (curvature: p < 0.001). In contrast, hit rate showed a significant but markedly smaller U-shaped relationship as a function of target frequency, indicating slightly reduced performance around the midpoint of the spectral distribution relative to the edges (curvature: p < 0.001; Fig. 4b). A direct comparison of curvature effect sizes revealed that, although both effects differed significantly from zero, adaptation to the distribution midpoint was substantially stronger for foreperiod duration than for target frequency (foreperiod and target frequency: p < 0.001; Fig. 4c). Reaction time showed a similar pattern. Reaction times were shorter around the midpoint of the foreperiod distribution than at its edges, reflected in a U-shaped relationship in the variable temporal condition (curvature: p < 0.001; Fig. 4d). Similar to hit rate, the modulation was asymmetric, with the fastest responses occurring at longer intermediate foreperiods rather than exactly at the distribution midpoint. To further characterize this asymmetry, we compared several PDF- and hazard-rate-based temporal-prediction models following Grabenhorst et al. (2019, 2026). This analysis is reported in Supplementary Fig. 2 and discussed below. In contrast to the foreperiod distribution, no significant curvature was observed for reaction time as a function of target frequency; reaction times remained stable across the spectral distribution (curvature: p = 0.629; Fig. 4e). This dissociation is also captured by the curvature effect size, which revealed a pronounced reaction time benefit around the foreperiod distribution midpoint but no reliable benefit around the midpoint of the target frequency distribution (foreperiod: p < 0.001; target frequency: p = 0.615; Fig. 4f).

**Figure 4-.**
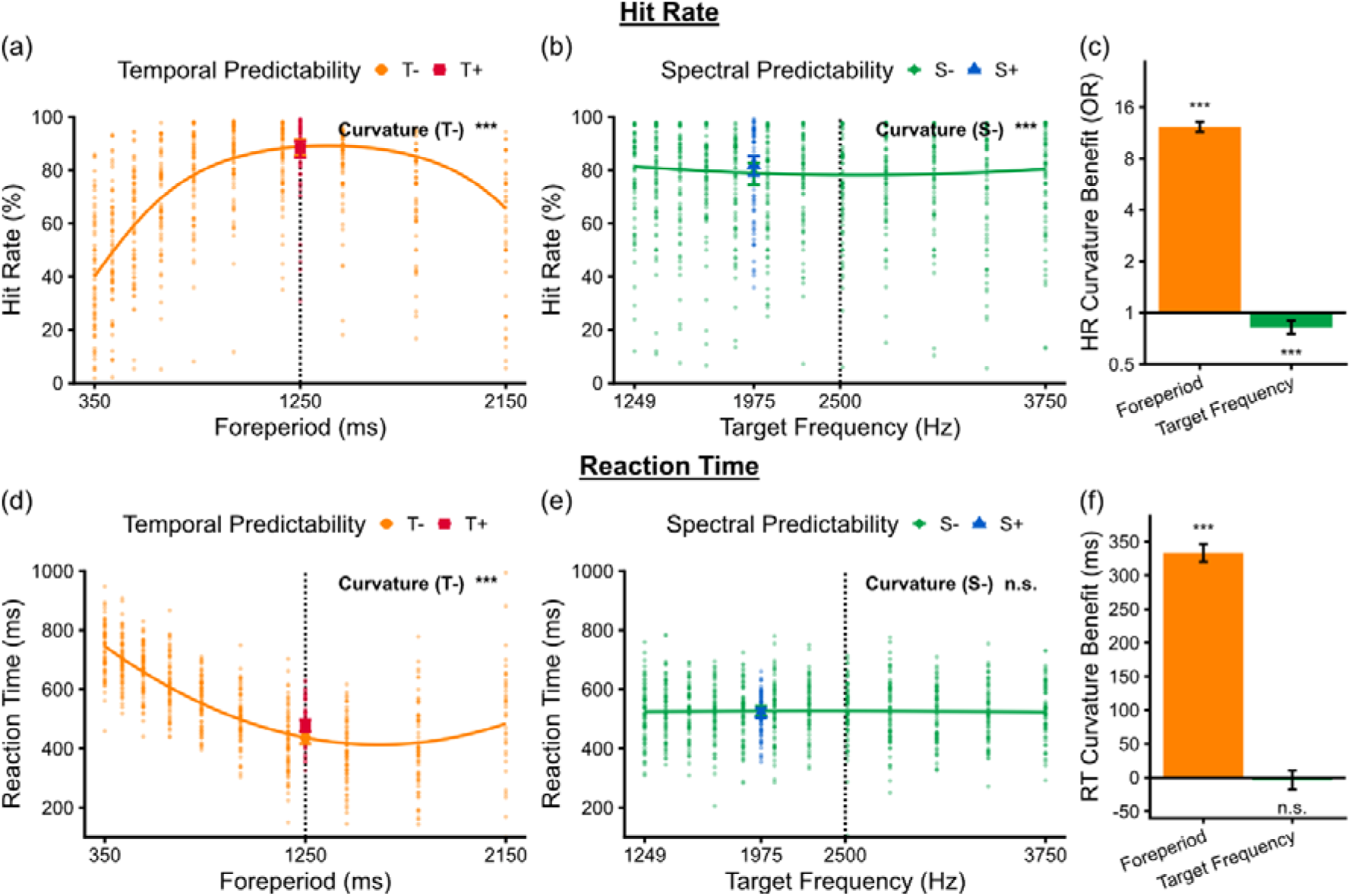
Distinct encoding strategies for temporal and spectral distributions. (a) Hit rate (%) as a function of foreperiod duration (ms; x-axis) and temporal predictability (color-coded). The solid line shows model-predicted performance across the variable foreperiod distribution (T-), and dots represent individual participant means. The vertical dotted line indicates the arithmetic midpoint of the foreperiod distribution. The label “Curvature (T-)” indicates a non-linear change in performance across the variable foreperiod distribution (T-); asterisks denote the statistical significance of this non-linear (quadratic) effect. Symbols show model-estimated marginal means for each condition at the fixed foreperiod duration, with whiskers indicating 95 % confidence intervals. (b) Same as panel (a), but showing hit rate (%) as a function of target frequency (Hz; x-axis) and spectral predictability (color-coded). The vertical dotted line indicates the arithmetic midpoint of the target-frequency distribution. (c) Effect size (OR) of the curvature effect on hit rate in the low-predictability condition. An OR greater than 1 indicates better performance around the midpoint of the distribution compared to its edges, whereas an OR smaller than 1 indicates worse performance around the midpoint relative to the edges. (d-f) Same as panels (a-c), but for reaction time.

A direct comparison of model-estimated performance at the fixed foreperiod and target-frequency values revealed no significant difference in hit rate between the fixed and variable temporal conditions at the fixed foreperiod duration (T-: HR = 88.80 ± 1.51 %; T+: HR = 88.17 ± 1.57 %; p = 0.074; Fig. 4a). In contrast, hit rate was slightly higher in the fixed spectral condition than in the variable spectral condition when evaluated at the fixed target frequency (S-: HR = 78.88 ± 2.10 %; S+: HR = 81.95 ± 1.86 %; p < 0.001; Fig. 4b). Reaction time showed a different pattern. When evaluated at the foreperiod used in the fixed temporal condition, reaction times were slower in the fixed than in the variable temporal condition (T-: RT = 436.58 ± 10.57 ms; T+: RT = 476.66 ± 10.15 ms; p < 0.001; Fig. 4d). For target frequency, reaction times were slightly faster in the fixed spectral condition compared to the variable spectral condition at the fixed frequency value (S-: RT = 526.13 ± 8.44 ms; S+: RT = 518.74 ± 8.26 ms; p = 0.025; Fig. 4e).

## 4. Discussion

### Complementary benefits of temporal and spectral predictions on auditory performance

The present results demonstrate that implicitly learned temporal and spectral predictions shape the perceptual and decisional processes underlying auditory detection in distinct ways. Temporal predictability alone increased response readiness: when target onsets were predictable, participants responded faster and produced more hits but also more false alarms, reflecting a liberalization of response bias at predicted moments in time. Spectral predictability, in contrast, selectively enhanced perceptual sensitivity, mainly by reducing false alarms, without producing a comparable speeding of responses. Combining both predictive dimensions yielded a synergistic increase in sensitivity but did not significantly affect response bias. The absence of a spectral predictability effect on reaction time differs from previous studies reporting that spectral predictions can influence response speed (Lange, 2008; Hsu et al., 2013). This discrepancy may reflect differences in task demands, task difficulty, or in how spectral expectations were induced.

The present findings reveal the distinct contributions of implicit temporal and spectral predictions to auditory detection and suggest how the auditory system may exploit predictive structure to improve perception and guide decisions under uncertainty: Temporal predictions direct preparation towards relevant moments in time, often described as temporal orienting of attention (Jones, 1976; Nobre and van Ede, 2018; Denison et al., 2021), reflected behaviorally in faster responses and a more liberal response bias. Spectral predictions, in contrast, prepare frequency-selective neural populations and thereby improve perceptual sensitivity, mainly through reduced false alarms (Okamoto et al., 2007; De Martino et al., 2015).

The dependence of temporal prediction benefits on spectral predictability replicates previous evidence (Hsu et al., 2013; Morillon et al., 2016; Wollman and Morillon, 2018) and is consistent with a hierarchical model of predictive filters, as proposed by Wollman and Morillon (2018). In this model, auditory processing is constrained by the tonotopic organization of the auditory system. Therefore, spectral predictions can be seen as a basic filtering level of auditory signals, whereas temporal predictions act at a higher level by enhancing an already selected frequency channel at the expected moment (Jaramillo and Zador, 2010; Lakatos et al., 2013; Fig. 1c). The present study extends this model by showing that the functional contribution of temporal predictions differs depending on the availability of spectral predictions. Without a prediction about the specific upcoming spectral content, temporal predictions exert a less selective effect, mainly increasing response readiness, as reflected in faster responses and a more liberal response bias. This is consistent with earlier reports of nonspecific gain modulation by temporal predictions (Auksztulewicz et al., 2018, 2019) and increased impulsive responding when stimulus timing is predictable (Korolczuk et al., 2018, 2020, 2022, 2023). When spectral predictions are available, however, temporal predictions can amplify feature-specific benefits and thereby further enhance perceptual sensitivity.

This interpretation also has implications for earlier studies in which temporal predictions appeared to improve sensitivity on their own. Such effects may have partly depended on interactions with implicit spectral predictability, particularly in paradigms using narrow frequency ranges (Lawrance et al., 2014; Herbst and Obleser, 2019). In these cases, specific temporal preparation may have been supported by feature-specific knowledge about upcoming stimuli, allowing the brain to prime the relevant sensory channels at the expected moment.

A similar division of labor between two predictive dimensions has been described in the visual domain, where predictions about stimulus probability and stimulus location exert separable effects on response processes: probabilistic cues primarily affect response bias, whereas location cues selectively enhance perceptual sensitivity (Bashinski and Bacharach, 1980; Reynolds et al., 2000; Carrasco, 2006; Wyart et al., 2012; Summerfield and de Lange, 2014). The present findings suggest that the conceptual principle of complementary predictive effects may generalize across sensory domains.

### Differential adaptation to temporal and spectral variability

Our findings demonstrate a qualitative dissociation in how implicit temporal and spectral variability shape behavior. When foreperiods varied, both hit rate and reaction time showed a pronounced center-weighted modulation, with higher hit rates and faster responses around the midpoint of the temporal distribution relative to its edges. Notably, both profiles were tilted towards longer intermediate foreperiods rather than being perfectly symmetric. This asymmetry suggests that temporal predictions may rely on more than one statistical property of the foreperiod distribution. To explore this possibility post-hoc, we compared PDF- and hazard-rate-based temporal prediction models following Grabenhorst et al. (2019, 2026; Supplementary Fig. 2). Across both reaction time and hit rate, the constantly-blurred hazard-rate model provided the best relative fit. This suggests that behavior at least partially reflected the hazard-rate (Nobre et al., 2007; Herbst et al., 2018). However, this model comparison should be interpreted cautiously because none of the tested models fully captured all aspects of the late-foreperiod pattern and the model comparison therefore does not provide a complete generative account of behavior. Additionally, the model ranking differs from Grabenhorst et al. (2019, 2026), where reciprocal PDF-based models outperformed hazard-rate-based models. One possible reason for this discrepancy is the different spacing of time points within the temporal distributions. Grabenhorst et al. used equally spaced time points across all distributions, while our foreperiods were logarithmically spaced. In our experiment, integration of temporal statistics across predictive and non-predictive blocks may also have contributed to the relatively good performance at intermediate foreperiods, as the intermediate foreperiod (1250 ms) was overall the most probable one. The observation that reaction times at 1250 ms were faster in the variable than in the fixed foreperiod condition was surprising. A possible explanation is that temporal precision decreases over longer fixed foreperiods, which can increase reaction times even when target timing is fully predictable (Niemi and Näätänen, 1981). Alternatively, high temporal predictability may have altered the temporal dynamics of sensory evidence integration, favoring more prolonged stimulus processing and consequently increased reaction times.

In contrast to the temporal distribution, performance across the spectral distribution showed no comparable midpoint-enhancement and remained largely stable. The same pattern was observed in a pilot experiment using a substantially narrower frequency range (1724–2224 Hz), suggesting that this near-flat spectral profile is largely invariant to the frequency range (Supplementary Fig. 1). This flat profile across target frequencies also argues against the possibility that the fixed cue frequency induced a strong frequency-repetition bias. One possibility is that frequency statistics were represented more veridically than temporal statistics: under a uniform distribution, optimal allocation of resources would yield relatively uniform performance across frequencies. Under this hypothesis, participants may have distributed attention relatively evenly across frequency channels in the variable condition and focused on a particular frequency in the predictive condition to enhance sensitivity. This interpretation is supported by recent work showing that auditory detection performance remains relatively uniform across equiprobable frequency distributions but becomes shaped by the distribution when frequencies differ in probability, with enhanced detection around high-probability frequencies and reduced detection of less-probable frequencies (Luthra et al., 2025).

## Conclusion

Our results reveal a simple organizing principle: temporal and spectral predictions jointly improve performance yet do so via complementary roles and distinct encoding strategies. Temporal predictions alone primarily modulate response readiness, whereas spectral predictions play an important role in selectively enhancing perceptual sensitivity at predicted moments. By dissociating these contributions, the present results refine our understanding of how the auditory system exploits and combines temporal and spectral statistics in the acoustic input to optimize perception under uncertainty.

## Author contributions

Study design: Sophie Herbst, Chloé Dumeige, Manon Beurtey, Johannes Wetekam

Experiments: Johannes Wetekam, Chloé Dumeige

Analysis and original draft of the manuscript: Johannes Wetekam, Sophie Herbst

Review and editing of the original draft: Sophie Herbst, Johannes Wetekam, Chloé Dumeige, Manon Beurtey

## Acknowledgements

This work was funded by the French National Research Agency (ANR-23-CE28-0006-01).

## Conflict of Interest

The authors declare no competing interests.

## Supplementary material

**Supplementary Figure 1-.**
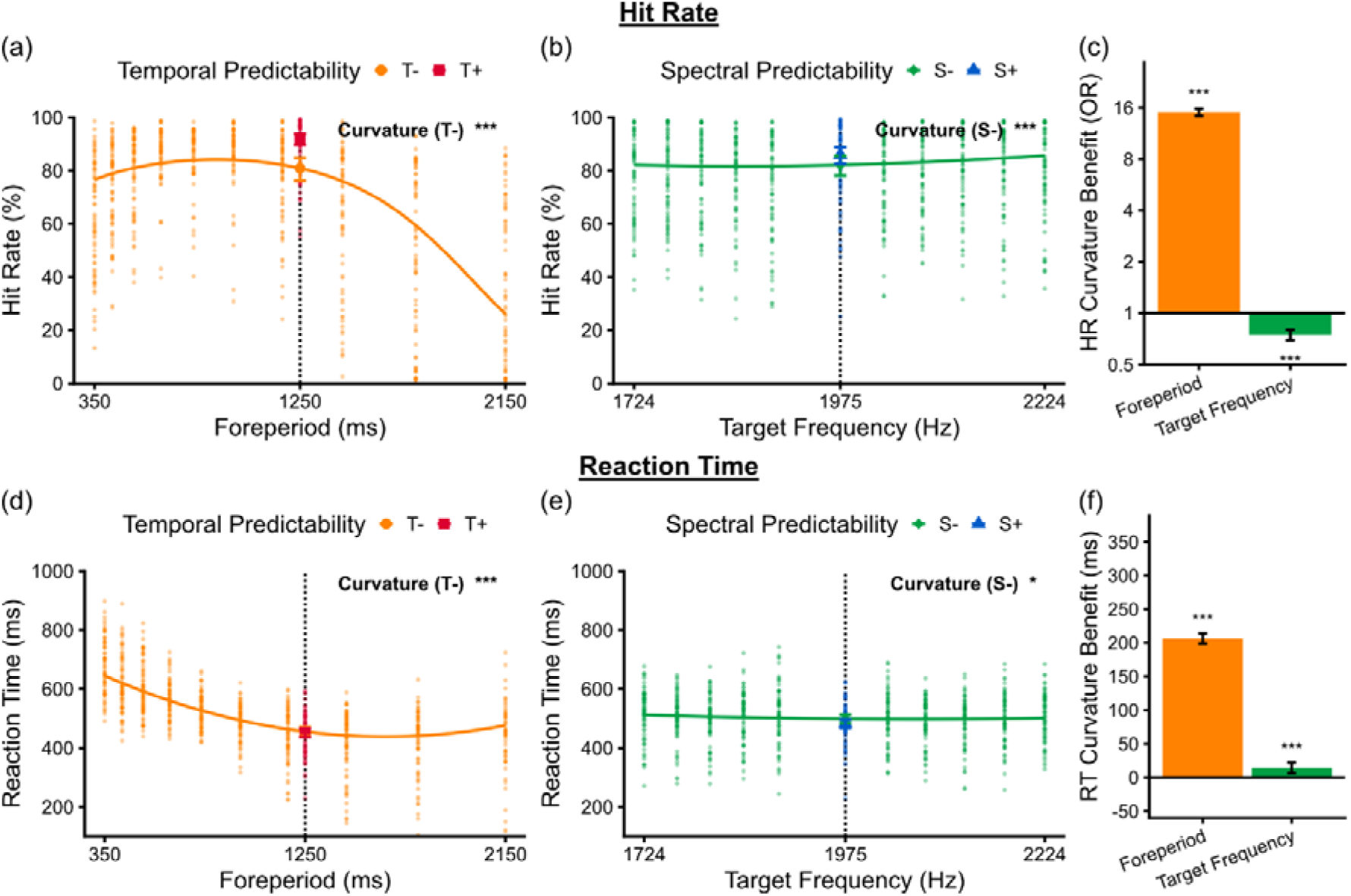
Distinct encoding strategies for temporal and spectral distributions in a pilot experiment with narrower frequency range. (a) Hit rate (%) as a function of foreperiod duration (ms; x-axis) and temporal predictability (color-coded). The solid line shows model-predicted performance across the variable foreperiod distribution (T-), and dots represent individual participant means. The vertical dotted line indicates the arithmetic midpoint of the foreperiod distribution. The label “Curvature (T-)” indicates a non-linear change in performance across the variable foreperiod distribution (T-); asterisks denote the statistical significance of this non-linear (quadratic) effect. Symbols show model-estimated marginal means for each condition at the fixed foreperiod duration, with whiskers indicating 95 % confidence intervals. (b) Same as panel (a), but showing hit rate (%) as a function of target frequency (Hz; x-axis) and spectral predictability (color-coded). (c) Effect size (OR) of the curvature effect on hit rate in the low-predictability condition. An OR greater than 1 indicates better performance around the centre of the distribution compared to its edges, whereas an OR smaller than 1 indicates worse performance around the centre relative to the edges. (d-f) Same as panels (a-c), but for reaction time.

**Supplementary Figure 2-.**
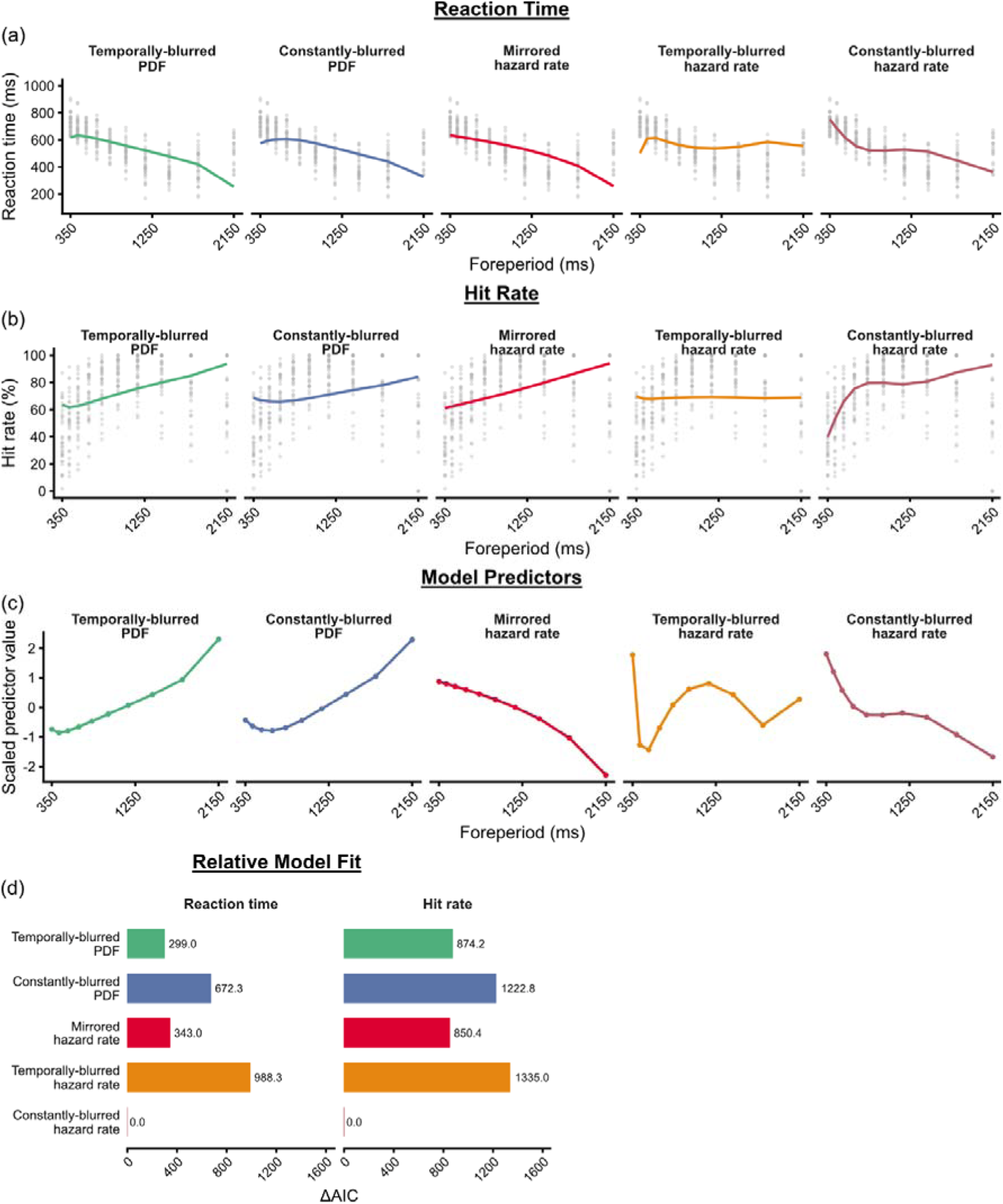
Comparison of PDF- and hazard-rate-based temporal-prediction models. (a) Reaction time (ms) as a function of foreperiod duration in the variable temporal condition. Grey dots represent individual participant means. Colored lines show fixed-effect predictions from single-predictor LMMs. (b) Same as panel (a), but for hit rate (%). Colored lines show fixed-effect predictions from single-predictor binomial GLMMs. (c) Scaled predictor values for the five tested temporal-prediction models. PDF predictors were entered as reciprocal PDF values, and hazard-rate predictors were mirrored, such that larger predictor values indicate weaker temporal expectation. (d) Model fit for reaction time and hit rate, expressed as ΔAIC, relative to the constantly-blurred hazard-rate model. Lower ΔAIC values indicate better relative model fit. Across both reaction time and hit rate, the constantly-blurred hazard-rate model provided the best relative fit.

## Notes

### Competing Interest Statement

The authors have declared no competing interest.

### Summary of Updates

We clarified the temporal model comparison and its interpretation, softened claims where evidence for a null effect was limited, revised the discussion of the hazard-rate effect, and added recent work relevant to the spectral-distribution results.

